# Tentorium cerebelli’s notch severity in the human brain: protocol, distribution and histopathology validation

**DOI:** 10.1101/2025.09.18.677182

**Authors:** Smriti Chadha, Joshua L. Chase, Emma W. Rosenblum, Erin-Marie C. Smith, Nathaniel Mercaldo, Matthew P. Frosch, Jean C. Augustinack

## Abstract

The dura forms an impression, the tentorial notch (TN), on the surface of the entorhinal cortex (EC). To characterize the potential injury to the EC, we evaluated the entorhinal surface in 55 postmortem samples. We developed a semi-quantitative protocol to rate the TN severity and evaluated the impact of the TN on the surrounding tissue. We rated the penumbral injury as: none, moderate or severe. We demonstrate that 96% of the samples showed a TN; specifically mild (33%), moderate (47%) and severe (16%). TN severity correlated positively with penumbra scores (Kendall’s coefficient = 0.65, 95% CI [0.535, 0.735]). We found no association between TN severity levels and Braak and Braak stages (Fischer’s exact *P* value = .862). We established a robust protocol for assessing the tentorial notch, addressing a gap in understanding anatomical factors that may contribute to EC vulnerability in Alzheimer’s disease.

## 1 Background

The entorhinal cortex (EC) exhibits early vulnerability in Alzheimer’s disease (AD), making it a critical region for understanding disease progression. While aging and genetic predisposition remain the most established risk factors for AD, understanding the selective vulnerability of the EC is essential from a molecular and anatomical perspective. Traumatic brain injuries have been linked as a risk factor for dementia[1–6]. The proximity, between the tentorium cerebelli (dura) and the entorhinal cortex may give rise to an injury, which is particularly relevant in conditions involving increased intracranial pressure. For example, in extreme cases of space-occupying lesions and elevated intracranial pressure, the tentorium cerebelli has been documented to compress and leave a severe nick on the parahippocampal gyrus, leading to transtentorial herniation[7–9]. This type of herniation leads to Duret hemorrhage of the brainstem, resulting in death[10].

Building on this concept of mechanical factors in the region, it is noteworthy that, even in individuals without any history of tumors or hemorrhages, the tentorium cerebelli creates a physical impression on the entorhinal cortex called the tentorial notch[11–14], sometimes referred to by uncal notch or intrarhinal sulcus[15–19]. The wear and tear exerted by the rigid tentorium against the cortical tissue may create localized areas of stress and vulnerability that could predispose these regions to inflammation, which may ultimately lead to neurodegeneration.

Given that the entorhinal cortex is the only region in close proximity to the free edge of the dura, and considering its early involvement in Alzheimer’s disease, it is essential to evaluate the effect of the indentation in greater detail. Therefore, in this study, using gross images, we develop a novel protocol to differentiate between the various severities of this indentation, hereby referred to as the tentorial notch. We also apply a semi-quantitative protocol to examine the impact of the tentorium cerebelli on the neurons in entorhinal cortex using Nissl stained coronal sections. We compare the tentorial notch severity rating to the Braak and Braak (BB) stages for each sample to evaluate whether TN severity may affect phosphorylated tau burden. This study provides a ground truth analysis around the impact of the tentorium cerebelli on the entorhinal cortex.

## 2 Methods

### 2.1 Sample demographic information

Fifty-five human brain hemispheres (n=27 left, n=28 right) were collected from the Neuropathology Autopsy Service at Massachusetts General Hospital, with approval from the Institutional Review Board. All hemispheres were fixed by immersion in 10% neutral buffered formalin for a minimum of 2 months and then transferred to 2% periodate-lysine-paraformaldehyde for storage at 4°C. The sample set included 23 females and 32 males between the ages of 20 and 84 years (median= 65, interquartile range [IQR] = 14). The postmortem interval for 35 cases was less than 24 hours; the other 20 samples had longer intervals due to the COVID-19 pandemic. We excluded samples with memory impairment (i.e. Alzheimer’s disease), Parkinson’s disease, stroke, or infectious diseases (e.g., HIV, hepatitis B, and prion disease). We also restricted the samples based on brain weight and only included samples greater than 1150 grams with no gross atrophy. We carefully screened for cortical atrophy not to confound the results. Table 1 summarizes the demographic information for this sample set.

**Table 1.** Demographic information for the *ex vivo* sample set. BB= Braak and Braak. g = grams, IQR = interquartile range.

| **Ex vivo Samples** | **Female**<br>***n*=23** | **Male**<br>***n*=32** | **Total**<br>***N*=55** |
| --- | --- | --- | --- |
| Age (years), median (IQR) | 60 (24) | 68 (14.5) | 65 (14) |
| Hemisphere *n* (%) |  |  |  |
| Left | 11 (41) | 16 (59) | 27 (49) |
| Right | 12 (43) | 16 (57) | 28 (51) |
| Brain weight (g), median (IQR) | 1288 (350) | 1344 (465) | 1325 (475) |
| BB Stage *n* |  |  |  |
| Normal Control | 7 | 11 | 18 |
| BBI | 2 | 4 | 6 |
| BBII | 3 | 6 | 9 |
| BBIII | 1 | 4 | 5 |

### 2.2 Entorhinal surface landmarks

The entorhinal cortex is surrounded by the rhinal sulcus anteriorly and the collateral sulcus laterally [20–26]. These sulci serve as important landmarks for identifying the entorhinal cortex. In addition to these sulci, the medial entorhinal cortex is characterized by the presence of the bulging structure known as gyrus ambiens (Brodmann’s area 34) [15]. Brodmann’s area 34 also corresponds to Insausti’s entorhinal intermediate medial subfield (EMI) [19,27]. Just inferior and lateral to the gyrus ambiens, an indentation by the tentorium cerebelli exists, which has been referred to as the tentorial notch [26,28,29] or the uncal notch [11,30,31]. Furthermore, a rare small shallow sulcus may be present within the entorhinal surface, which Heinsen referred to as the entorhinal sulcus [32,33]. We used these landmarks to accurately assess the location of the tentorial notch and its underlying cortex.

### 2.3 Reconstruction of Visible Human Project^®^ Data

We used the Visible Human Project^®^ data 2.0 [34] to analyze the proximity of the tentorium cerebelli to the entorhinal cortex (Figure 1). The specimen was a 72-year-old male, fixed with formalin. The data from the Visible Human Project^®^ can be accessed on the NIH website (https://www.nlm.nih.gov/research/visible/visible_human.html). The Visible Human Project^®^ data were originally collected in axial slices, made from an axial series of photographed blockface images. The images were also reconstructed to produce images in the coronal plane. We annotated the coronal images to show the dura’s point of contact with the entorhinal cortex (Figure 1A, unannotated and Fig. 1B annotated). The Visible Human Project^®^ brain included injection of the blood vessels (red for arteries; blue for veins), providing multiple landmarks. Figure 1 (A and B) illustrate the indentation, also known as the tentorial notch, and the proximity of tentorium cerebelli to entorhinal cortex. The Visible Human Project^®^ data allowed us to investigate and show all structures, not just soft tissue as in MRI. Further, in MRI, the medial temporal lobe suffers from B_0_-related signal dropout and distortion in this region, making the Visible Human Project^®^ data ideal to show dura, brain tissue and landmark vessels.

**Figure 1.**
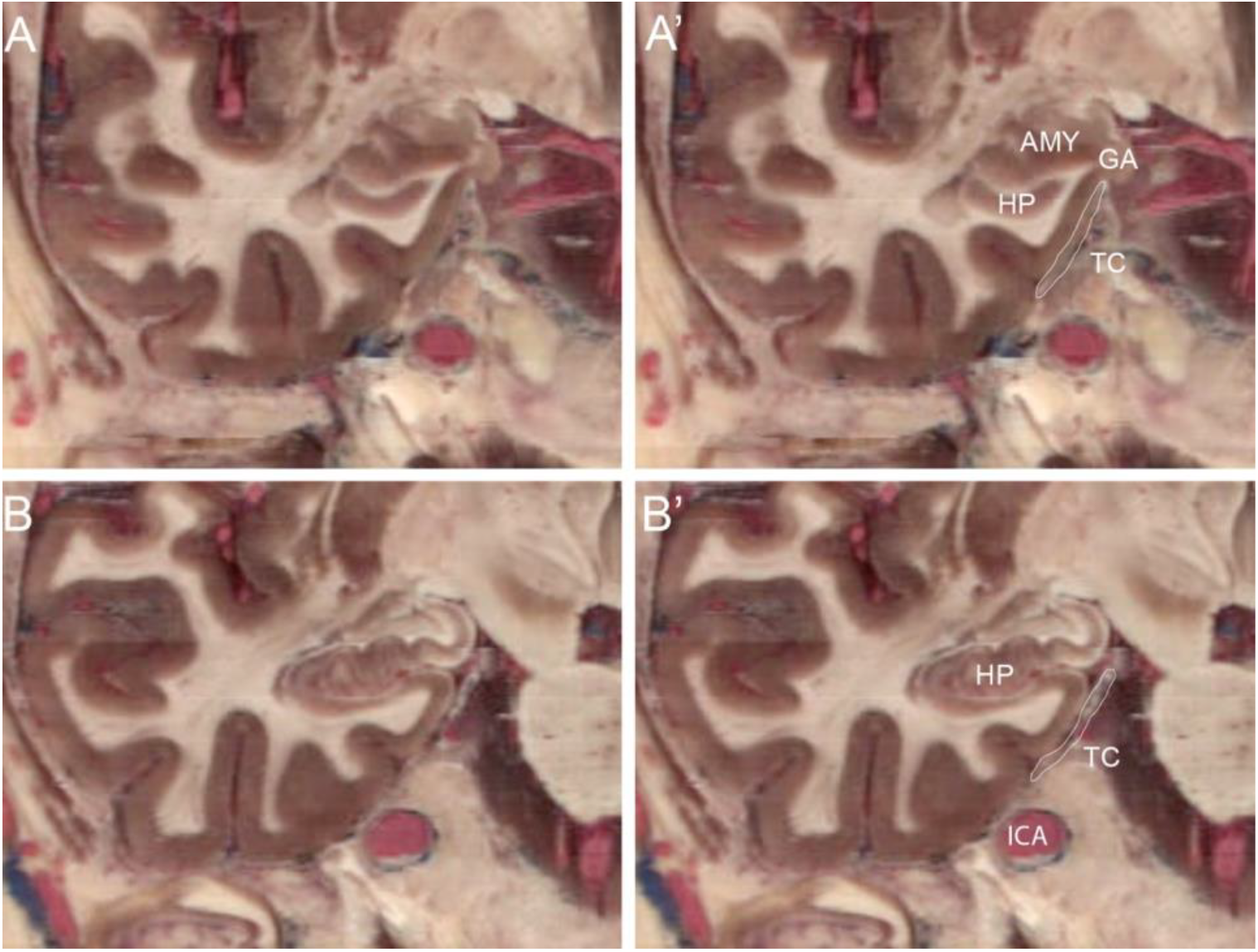
Reconstructed Medial Temporal Lobe Images from the Human Visibility Project. Panels A and B show the medial temporal lobe in the coronal plane at an anterior level of entorhinal cortex near amygdala and a posterior level of entorhinal cortex (level of hippocampal pes), respectively. The same images with annotations are represented in panels A’ and B’. Note the proximity of the free edge of the tentorium cerebelli to the gyrus ambiens (Brodmann’s GA or Insausti’s EMI in A’ but not B’. White outline denotes the tentorium cerebelli in A’ and B’. Abbreviations: AMY = Amygdala, GA = Gyrus Ambiens, HP = Hippocampus, ICA = Internal carotid artery, TC= Tentorium Cerebelli.

### 2.4 Photography of samples

We photographed each sample to catalog the data. Leptomeninges and brainstem were removed from each sample before photography to ensure clear visualization of the entorhinal cortex. Each medial temporal lobe was placed in a ventromedial position and photographed using a digital Canon PowerShot SX520 HS camera. The photographs were used to evaluate the entorhinal cortex surface to differentiate between the tentorial notch and entorhinal sulcus. The severity (depth) of the tentorial notch was evaluated on the photographs using the novel protocol described below.

### 2.5 Tentorial notch severity protocol

To differentiate between the severity levels of the tentorial notch, we developed a semi-quantitative rating scale to classify the depth of the indentation on the surface of the entorhinal cortex, using scores ranging from 0 to 3 (Figure 2A-D). *Score 0* (Figure 2A) denoted the absence of a tentorial notch. *Score 1* indicated a slight (mild) indent formed on the entorhinal surface (Figure 2B). *Score 2* represented a moderate notch impression (Figure 2C), whereas *score 3* designated a severe impression on the entorhinal surface (Figure 2D). Two raters (SC and JCA) evaluated and rated the severity of each tentorial notch using the ventromedial photographs.

**Figure 2.**
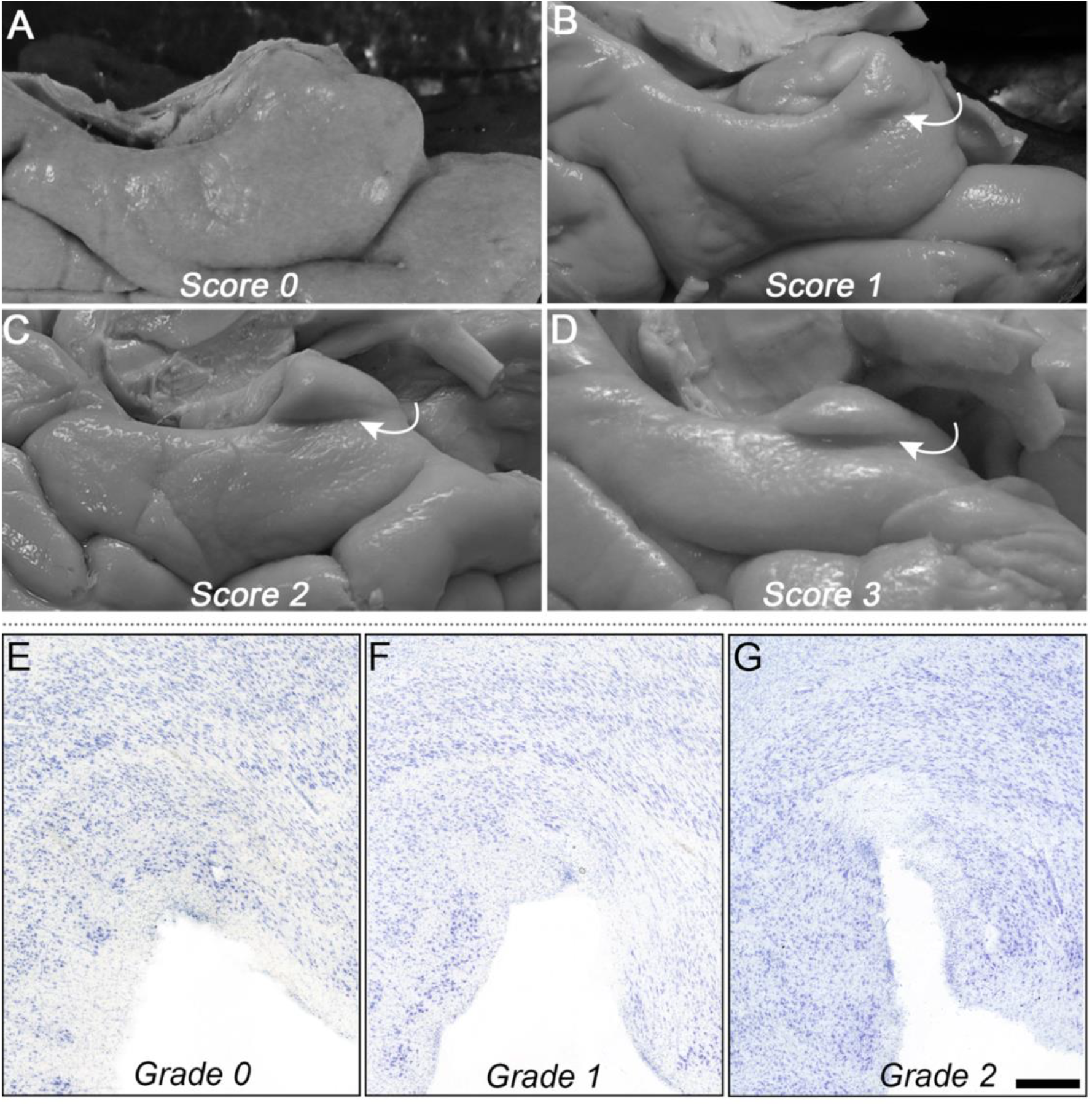
Semi-quantitative protocols for tentorial notch severity scores and penumbra grades. Panels A-D show the scoring for TN on the gross surface of entorhinal cortex, Tentorial notch severity scores: A) *Score 0*: lack of a tentorial notch. B) *Score 1*: shows a slight imprint (notch) inferior to the Gyrus Ambiens C) *Score 2:* a moderate notch. D) *Score 3:* a severe indentation is present. White arrows point to the tentorial notch. Panels E-G show the penumbra grading scores for Nissl histology. E) *Grade 0*: not impacted, typical neurons and layers. F) *Grade 1*: region shows neuronal shrinkage and evident disruption of layers I, II and III, slight penumbra. F) *Grade 2:* shows a severe penumbra with severe neuronal loss in layers II and III. Magnification bar = 1mm.

### 2.6 Tissue processing

For a subset (n=33) of cases, selected based on availability of tissue, hemispheres were blocked to section the underlying cortex. The blocks were placed in cryoprotectant (20% glycerol, 2% dimethyl sulfoxide) for a minimum of 2 months. Following cryoprotection, the blocks were sectioned serially at 50 µm on a freezing sliding microtome (Leica SM2000R or SM2010R, Leica Biosystems Inc.). Blocks were sectioned in the coronal plane. Blockface images of each section were collected using a mounted Canon EOS-1D Mark IV camera (Canon, Tokyo, Japan). All sections were stored in cryoprotectant at −20°C until further processing. Tissue sections for these cases were used in Nissl staining and immunohistochemistry as described below.

### 2.7 Nissl staining

Using blockface images, three tissue sections where the TN was present was chosen per sample. The tissue sections were thawed and rinsed in 0.1 M phosphate buffer prior to being mounted on gelatin-subbed slides and dried overnight. We followed a previously published Nissl protocol [35]. To defat the tissue sections, slides were placed for 20 minutes in 1:1 chloroform and 100% ethanol followed by three minutes each in 50% ethanol and double distilled water (ddH_2_O). Tissue was then pretreated in 1:1:1:1 glacial acetic acid, 100% ethanol, ddH_2_O, and acetone for one minute, followed by a one-minute wash in ddH_2_O. A buffered thionin solution (5% thionin, 1.36% sodium acetate stock, and 0.6% acetic acid stock) was used to stain the tissue for five minutes. Tissue was differentiated in 70% ethanol with a few drops of glacial acetic acid and dehydrated in a series of increasing ethanol solutions (70%, 95%, and 100%). Dehydrated slides were then transferred into xylene and coverslipped with Permount (Fisher Scientific, Waltham, MA).

### 2.8 Immunohistochemistry

Immunohistochemistry was performed according to the protocol described in [36]. CP13 is a mouse monoclonal antibody at serine 202 in tau. Briefly, cryoprotectant was rinsed from free-floating tissue using 1X phosphate-buffered saline. The sections were quenched using 0.5% Triton X-100 and 3% hydrogen peroxide for 20 minutes. Nonfat milk (5% in 1X PBS) was used to block non-specific binding at room temperature for 2 hours and sections were subsequently incubated for 48 hours at 4°C in primary antibody (CP13, Feinstein Institute for Medical Research, NY) at 1:200 dilution in 1.5% normal goat serum. An HRP-based mouse secondary was used at a 1:200 dilution in 1.5% normal goat serum for 1 hour. Chromogen was visualized with 3’3-diaminobenzidine (DAB kit, Vector Laboratories, Burlingame, CA). The tissue sections were rinsed and mounted on subbed gelatin slides and then dried overnight at room temperature. Defatting and dehydration steps from the histochemistry protocol (above) were applied, slides were cleared in xylene and coverslipped with Permount. The immunohistochemistry slides were used to stage samples according to Braak and Braak protocol [37]. We also used the immunohistochemistry slides to observe whether pTau levels correlated with tentorial notch severity ratings.

### 2.9 Penumbra grade protocol

To further examine the impact of the tentorium cerebelli on the entorhinal cortex, we evaluated histologically stained tissue immediately surrounding the tentorial notch on the Nissl stained sections. To differentiate between the variability of the impact, we developed a semi-quantitative grading scale as shown in Figure 2 panels E-G. *Grade 0* (Figure 2E) represents no injury. *Grade 1* (Figure 2F) exhibits moderate disruption in layers I, II and III in the entorhinal cortex along with neuronal atrophy. There was also a slight penumbra (white space depicting a loss of neurons) present. *Grade 2* (Figure 2G) illustrates a severe disruption in layers I, II and III that was observed, along with a severe penumbra depicting neuronal loss in layers II and III. Two raters evaluated and rated the impact on three Nissl sections in 33 samples.

### 2.10 Statistics

Descriptive summaries were computed for the entire cohort. Continuous variables were summarized through the median and interquartile range (IQR; 25-75 percentiles), whereas the categorical variables were characterized as frequencies (percentages). Gwet’s AC2 [38] was performed to analyze the inter-rater reliability for the tentorial notch severity protocol and the penumbra grading protocol. Ordinal logistic regression modeling was performed to look at the odds ratio between tentorial notch severity rating and sex. Since there were three sections per sample for Nissl, it was specified in the code to randomly sample one slide per case and report Kendall’s tau median and 95% confidence interval (CI) to measure the correlation between the tentorial notch severity and the penumbra scores. We also performed a Fischer’s exact test to determine an association between TN severity and Braak and Braak (BB) stages. All statistics were performed using R (version 4.4.1).

## 3 Results

### 3.1 Tentorial notch severity distribution

We observed that the tentorial notch was present in 96% of the samples (n=53/55 samples). The tentorial notch was further assessed with a semiquantitative rating (Figure 2) based on the severity of the imprint. Table 2 summarizes the distribution of the tentorial notch severity by sex and hemisphere. Briefly, tentorial notch was present as mild in 33% (n=18), moderate in 47% (n=26) and severe in 16% (n=9) of the samples. Gwet’s AC2 showed 94% agreement between rater 1 and 2, indicating protocol reliability as 80-100%, which indicates almost perfect agreement [39]. Rater 1 scores were used for all other statistics. Applying an ordinal logistic regression, we observed that females were 1.38 times more likely than males to exhibit a severe tentorial notch; however, this was not significant (*P* value = .53).

**Table 2.** Distribution of the tentorial notch and entorhinal sulcus according to sex and hemisphere.

| Landmarks | Frequency *n* (%) |  |  |  |
| --- | --- | --- | --- | --- |
|  | Sex |  | Hemisphere |  |
|  | Female<br>(n=23) | Male<br>(n=32) | Left<br>(n=27) | Right<br>(n=28) |
| **Tentorial notch severity** |  |  |  |  |
| None | 0 (0) | 2 (6) | 0 (0) | 2 (7) |
| Mild | 9 (39) | 9 (28) | 10 (37) | 8 (29) |
| Moderate | 8 (35) | 18 (56) | 15 (56) | 11 (39) |
| Severe | 6 (26) | 3 (10) | 2 (7) | 7 (25) |
| **Entorhinal sulcus** |  |  |  |  |
| Absent | 22 (96) | 25 (78) | 24 (89) | 23 (82) |
| Present | 1 (4) | 7 (22) | 3 (11) | 5 (18) |

### 3.2 Penumbra scores

Figure 2E-G demonstrate the rating scale used to characterize the impact of the tentorium cerebelli damage. We observed that *grade 0* (Figure 2E), no penumbra, was observed in 12 samples (36%) whereas *grade 1* (Figure 2F), indicating a moderate penumbra was observed in 39% of the samples. A severe penumbra, *grade 3* (Figure 2G) was observed in 24% of the samples. Gwet’s AC2 showed a 83% agreement between rater 1 and rater 2, indicating protocol reliability as 80-100%, indicating almost perfect agreement [39]. Kendall’s tau showed a median tau coefficient of 0.644 (95% CI [0.535, 0.735]) indicating substantial positive correlation between tentorial notch severity and the penumbra scores. Table 3 shows the correlation matrix between the tentorial notch ratings and the penumbra scores for all three sections for 33 samples.

### 3.3 Tentorial notch with underlying tissue architecture

Figure 3 illustrates multiple photographs showing the variability of tentorial notches whereas Figure 4 A-D displays the absence and three respective tentorial notch severities in stained Nissl sections that complement the gross photographs. In sections with a tentorial notch present (Figure 4B-D), we observed that subfield EMI was present medial to the tentorial notch whereas subfields ER and EI were present on the lateral portion. Figure 4D illustrates *score 3* and an especially deep indentation, denoting a severe tentorial notch in tissue.

**Figure 3.**
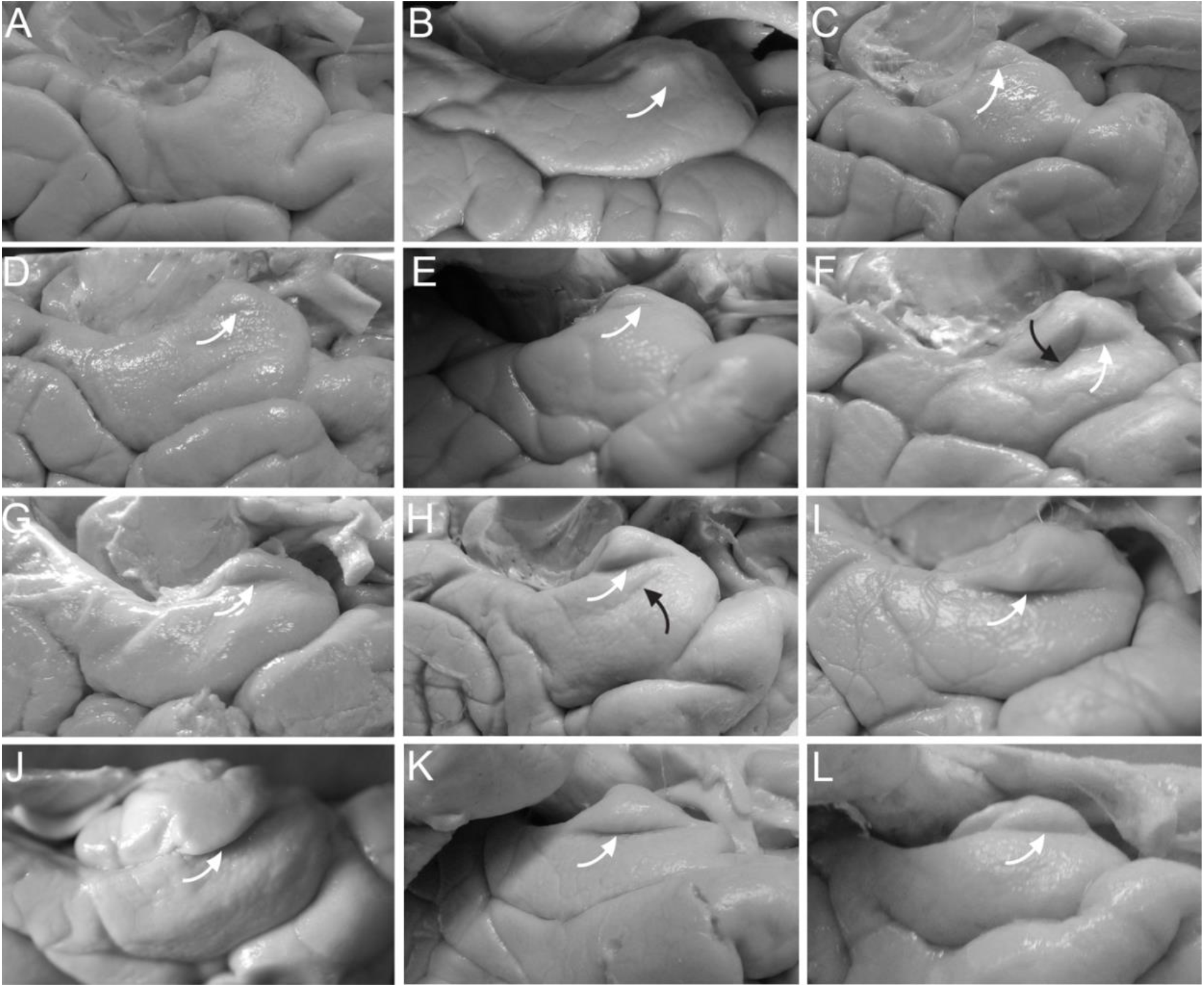
Variability in Tentorial Notch and Neighboring Structures. Panels A–L illustrate differences in the tentorial notch appearance and severity. Panel A displays an example of the lack of a tentorial notch. Panels B–D show mild notches (*Score 1*), panels E–H depict moderate notches (*Score 2*), and panels I–L demonstrate deep, pronounced notches (*Score 3*). White arrow indicates the tentorial notches, while black arrows in panel F and H indicates the entorhinal sulcus.

**Figure 4.**
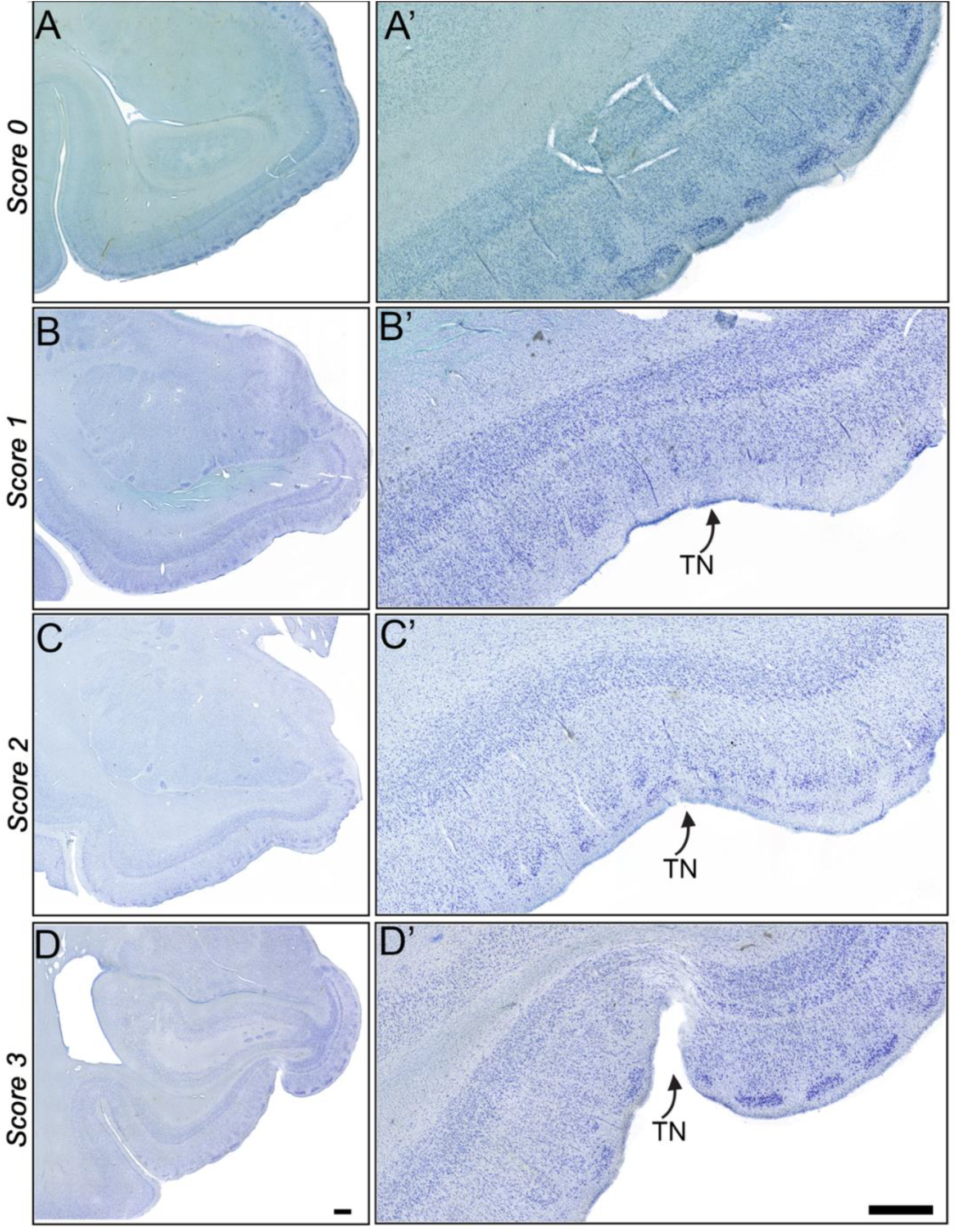
Tentorial notch severities illustrated in histology. Nissl stained sections in A-D. Panel A shows the lack of tentorial notch (*score 0*). Panels B-D depict the mild (*score 1*), moderate (*score 2*) and severe (*score 3*) tentorial notch, respectively. Panels A’-D’ show an enlarged view around the tentorial notch region from Panels A-D. The gyrus ambiens (of Brodmann) or Insausti’s EMI) was present on the medial side of the tentorial notch. Note a severe penumbra in D’. Panels A and D correspond to the same samples as in Fig. 3A and 3K, respectively and B and C show the same samples as in Fig. 2B and 3C, respectively. Magnification bar = 1 mm.

### 3.4 Entorhinal sulcus

An entorhinal sulcus was observed in 15% of the samples (n=8/55). Figure 3H and F shows the entorhinal sulcus on the gross image of two different samples. The entorhinal sulcus was present in just one female sample (4% of females) and seven male samples (22% of males). Table 2 summarizes the distribution of entorhinal sulcus according to sex and hemisphere.

### 3.5 pTau and TN severity

Figures 5A to F show the pTau reactivity in an anterior section at the level of TN (A, C, E) and a posterior section (B, D, F) entorhinal cortex in three different samples. We observe that the reactivity of pTau surrounding the tentorial notch is less than the reactivity in the lateral and posterior areas (in the same section). This implies that pTau reactivity does not begin near the tentorial notch. A Fischer’s exact test showed no association between BB staging and TN severity levels (*P* value = .862).

**Figure 5.**
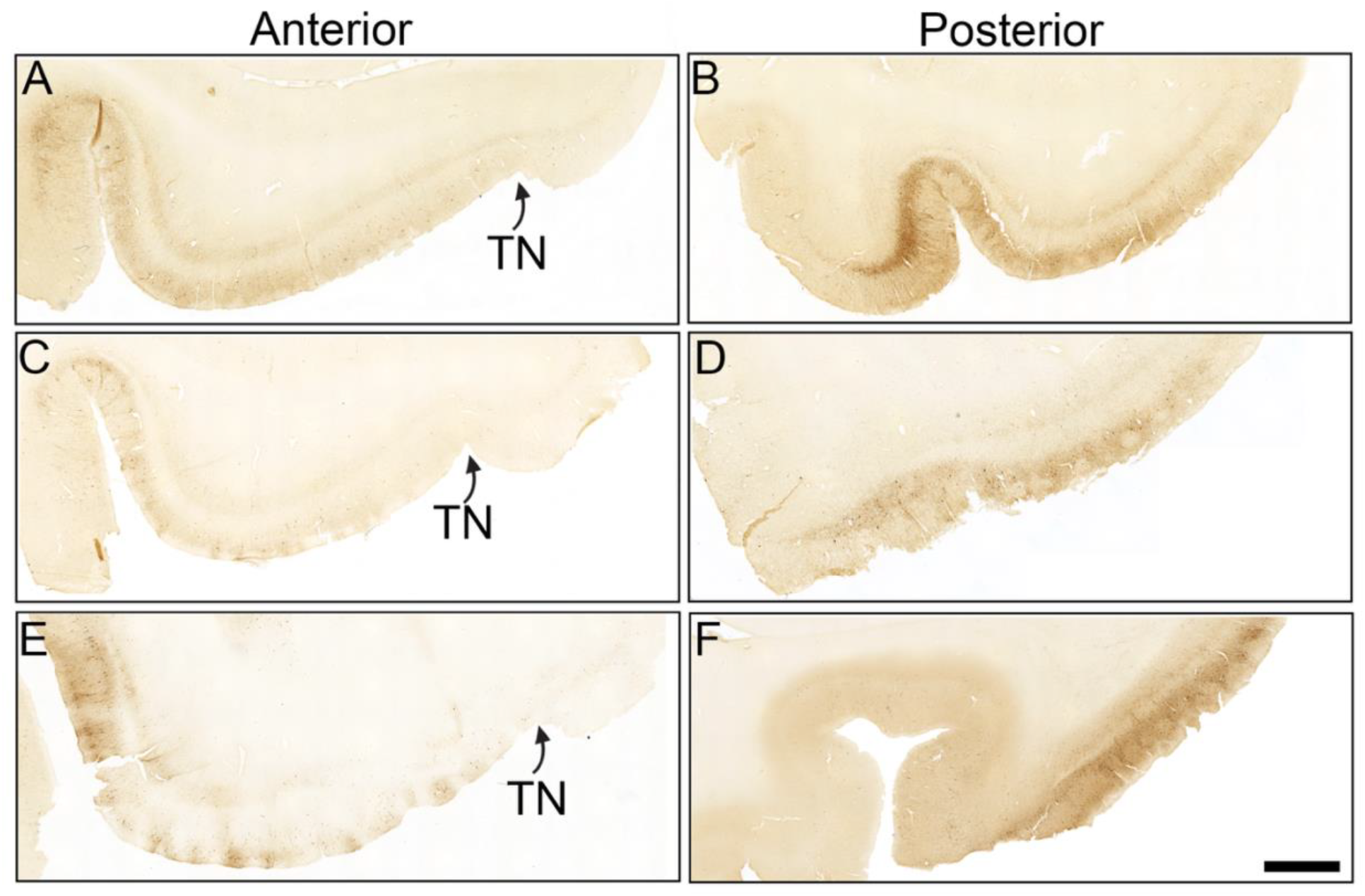
pTau in anterior and posterior entorhinal cortex of three samples. Panel A, C and E display the phosphorylated tau immunoreactivity in anterior entorhinal cortex. Panel B, D and F depict the phosphorylated tau immunoreactivity in posteriorly and medially in entorhinal cortex of the same cases as A, C and E, respectively. Higher ptau reactivity is observed laterally compared to medial entorhinal cortex in both anterior and posterior sections. More phosphorylated tau is observed in the posterior sections compared to anterior sections where the tentorial notch is present. Note the tau pathology is not at the location of the tentorial notch. All tissue sections are oriented such that the lateral portion of the EC is on the left and the medial is on the right. Magnification bar = 1mm.

## 4 Discussion

This study aimed to document the severity and distribution of the tentorial notch while assessing the impact of this injury on the entorhinal cortex tissue. We evaluated the gross surface of the entorhinal cortex to differentiate between the various severities of the tentorial notch. We also looked at the surrounding tissue to assess the impact of the tentorium cerebelli on the entorhinal cortex. Our observations revealed that the tentorial notch was present in 96% of our cases and is thus an extremely common occurrence in the human brain. Most postmortem samples showed a moderate tentorial notch (n = 26/55) whereas mild and severe notches were present in n = 18 and n = 9 samples respectively. Notably, all samples, including those rated as moderate or severe, had no history of herniation, brain tumors, or increased intracranial pressure. We also observed that no penumbra, was observed in 36% whereas *grade 1* was observed in 39% of the samples. A severe penumbra, *grade 3* was observed in 24% of the samples. There was no significant correlation between pTau severity and tentorial notch severity ratings. This may suggest that the close proximity of the tentorium cerebelli (and resultant notch) may represent an area of further study to investigate another variable, risk factor, or at least a vulnerability to disease.

The dura folds into the falx cerebri, falx cerebelli, diaphragma sellae, and tentorium cerebelli to protect and compartmentalize various brain regions. Due to the location, the free edge of the tentorium cerebelli is in close contact with the entorhinal cortex (Figure 1) in the human brain [11,12,14,28] and this proximity can lead to herniation in cases with increased intracranial pressure [7–9]. The close contact between these structures can also cause subclinical damage that is understudied. The severe tentorial impression observed in Figure 4D illustrates how a severe notch in this region may create a disruptive wear and tear injury and cause layer disruption. A severe penumbra was present in 24% of the samples. It has been previously shown that following traumatic brain injury, there is an infiltration of lymphocytes accompanied by microglial activation, which can lead to neurodegeneration [40–43]. The close contact between dura and EC may compromise tissue integrity, which could speculatively affect the blood brain barrier and initiate an inflammation cascade. Since tau phosphorylation in the EC initially accumulates in the lateral subfields rather than EMI and ER (where the TN is located), it is pertinent to study the different pathways through which the TN may contribute to entorhinal cortex vulnerability.

The term tentorial notch has been used before to describe the space between the free edges of the tentorium cerebelli [44] but based on Van Hoesen 1999[28], we believe the term tentorial notch is better suited for the indentation formed on the surface of the entorhinal cortex. While the nomenclature used to describe this structure may vary across studies, the distribution of the tentorial notch (indentation) observed in our sample aligns with several prior reports. Specifically, [19,33,45] all reported that the tentorial notch was present in nearly all cases examined. In contrast, our findings differ from those of Hanke (1997) and Retzius (1896), who reported the presence of the tentorial notch in only 69.6% and 54% of cases, respectively. The discrepancy may be attributed to subtle morphological variations—such as the notch presenting as a slight indentation— that could lead to underreporting or misidentification in gross anatomical assessments.

We also corroborated the presence of a rare sulcus, known as the entorhinal sulcus on the surface of the entorhinal cortex [13,19,32,33]. In our study of 55 samples, the entorhinal sulcus was observed in only eight cases. Heinsen previously reported the presence of the entorhinal sulcus in approximately 4% of cases and we report a 15% occurrence. Our data on the entorhinal sulcus shows a sex difference, consistent with previous finding by Heinsen[33] who reported that in 17 samples with an entorhinal sulcus, only 4 samples were females. The entorhinal sulcus was observed inferior to the tentorial notch. It is important to distinguish between these two because the tentorial notch may cause damage but the entorhinal sulcus is a naturally occurring sulcus. The entorhinal cortex varies slightly in size and shape [46–48] but the subfields ER and EI were consistently lateral to the tentorial notch. The characterization of this area – including the Visible Human Project^®^ data, ventromedial photographs and histology staining – recognizes and shows that the tentorial notch is almost always formed inferior to gyrus ambiens, between Brodmann areas 34 and 28.

It is well documented that females exhibit a higher vulnerability to Alzheimer’s disease and not just because females live longer than males [49–52]. In our data, we observed that women were 1.38 times more likely to have a more severe tentorial notch. While the results were not statistically significant, which may be due to a low n for a population study, there was a trend observed. Given high female risk to Alzheimer’s disease, it seems prudent and pertinent to investigate any risk factors that may be another piece of the puzzle. Future longitudinal studies may provide more information. While our paper evaluates the effect of this notch, it has previously been shown in MRI that the close proximity of dura to the entorhinal cortex [53,54] may be hard to quantify. Standard MRI protocols for brain morphometry exhibit B_0_-related signal distortions in this region, and state-of-the-art automated brain segmentation algorithms are not optimized to accurately segment the relevant tissues from these images. Further improvement of these protocols and methods would contribute to the study of dura proximity.

By evaluating the underlying tissue impacted by the tentorium cerebelli, we observed that there was a variability in the impact of the injury on the entorhinal cortex (Figure 5A-C). This injury was characterized into none, moderate and severe penumbra based on the disruption of the layers in entorhinal cortex along with neuronal atrophy or loss (Figure 2E-G). Notably, there was a positive correlation between the severity of the tentorial notch and the penumbra scores (Kendall’s tau median = 0.65) but no significant correlation between TN severity and the Braak and Braak stages (Fischer’s exact *P* value = .862). Thus, our study suggests that the tentorial notch plays a role in causing layer disruption and neuronal loss in entorhinal cortex but it may not play an initial role in tau phosphorylation. It has been previously shown that after locus coeruleus, tau accumulation occurs in perirhinal cortex along with posterior and lateral entorhinal subfields [37,55–59] rather than at the site of the tentorial notch near subfields EMI and ER. It could be speculated that this region of neuronal loss and layer disruption may act as a triggering factor for inflammation. Contrarily, the tentorium cerebelli is closer to the entorhinal cortex anteriorly and medially, showing that this injury likely does not contribute to tau accumulation.

Our study had some limitations. Our samples were collected from a hospital population; therefore, the findings may not represent a community population. We did not have access to both hemispheres from the same individual; thus, the interhemispheric variability of the tentorial notch and entorhinal sulcus was not studied. Finally, the nature of the study was cross-sectional, not longitudinal, and we were not able to assess trajectories of the tentorial notch and its impact. Most samples were aged but we excluded samples with low weight, gross atrophy and neurodegenerative diseases. Future studies could examine the effect of aging on TN severity. Our study also had many strengths. Compared to other anatomical studies, we had a large sample set and we established a novel protocol regarding the tentorial notch severity and its impact on the entorhinal cortex. We assessed on multiple histology and histopathology sections. The detailed gross anatomical and cytoarchitectural analyses on the same brain samples provide a macroscopic and microscopic scale examination of the tentorial notch. We selected cases with no known history of dementia which allowed us to focus on the impact of the tentorial notch without including confounding variables like neuronal loss from dementia. This integrated methodology provides a more precise and biologically grounded framework.

In conclusion, this study not only serves as an observational study but also establishes two novel protocols for the brain and dura relationship by providing a broader picture of the central nervous system environment. We show that while the tentorial notch may resemble tissue damage and disrupt entorhinal cortex layers, the mechanism or long term effect is not known. Here, we provide a foundational lens that subsequent studies may build upon to understand the mechanisms involved in Alzheimer’s disease progression.

## Acknowledgements

We thank our brain donors and their families who made this work possible. We also thank Dr. André v an der Kouwe for the thoughtful discussion. We acknowledge the late Dr. Peter Davies’s generosity for the CP13 tau antibody.

## Consent Statement

Autopsy consent was obtained, and tissue was collected with allowance for tissue to be used in research under a protocol approved by the Institutional Review Board of Mass General Brigham.

## Data Availability Statement

All supporting data are available in the paper and Supplementary Materials.

